# Assessing the ability of ChatGPT to extract natural product bioactivity and biosynthesis data from publications

**DOI:** 10.1101/2024.08.01.606186

**Authors:** Thomas L. Kalmer, Christine Mae F. Ancajas, Zihao Cheng, Abiodun S. Oyedele, Hunter L. Davis, Allison S. Walker

**Affiliations:** Department of Chemistry, Vanderbilt University, Nashville, TN USA; Department of Biological Sciences, Vanderbilt University, Nashville, TN USA; Department of Pathology, Microbiology, and Immunology, Vanderbilt University Medical Center, Nashville, TN, USA

## Abstract

Natural products are an excellent source of therapeutics and are often discovered through the process of genome mining, where genomes are analyzed by bioinformatic tools to determine if they have the biosynthetic capacity to produce novel or active compounds. Recently, several tools have been reported for predicting natural product bioactivities from the sequence of the biosynthetic gene clusters that produce them. These tools have the potential to accelerate the rate of natural product drug discovery by enabling the prioritization of novel biosynthetic gene clusters that are more likely to produce compounds with therapeutically relevant bioactivities. However, these tools are severely limited by a lack of training data, specifically data pairing biosynthetic gene clusters with activity labels for their products. There are many reports of natural product biosynthetic gene clusters and bioactivities in the literature that are not included in existing databases. Manual curation of these data is time consuming and inefficient. Recent developments in large language models and the chatbot interfaces built on top of them have enabled automatic data extraction from text, including scientific publications. We investigated how accurate ChatGPT is at extracting the necessary data for training models that predict natural product activity from biosynthetic gene clusters. We found that ChatGPT did well at determining if a paper described discovery of a natural product and extracting information about the product’s bioactivity. ChatGPT did not perform as well at extracting accession numbers for the biosynthetic gene cluster or producer’s genome although using an altered prompt improved accuracy.

## Introduction

Natural products, also referred to as secondary or specialized metabolites, are small molecules produced by living things and are distinct from those produced through primary metabolism. Natural products have played an important role in drug discovery. Many FDA approved drugs are natural products, natural product derivatives, or are synthetic compounds that inhibit a target originally identified through studies on a natural product.^1^ Early natural product discovery efforts relied mostly on screens of microbes from prolific taxa such as Actinomycetes or Penicillium. Over time, this strategy has become less successful as the most common natural products from prolific taxa have already been discovered, leading to frequent rediscovery of known compounds.^2^ Therefore, the field has increasingly moved to more targeted strategies for discovering novel bioactive natural products.

Genome mining, a strategy in which bioinformatic tools are used to assess an organism’s biosynthetic capacity, is one strategy to address the challenge of rediscovery. Bacterial, fungal, and some plant natural products are produced by a set of biosynthetic enzymes encoded by genes in close proximity in the genome, referred to as a biosynthetic gene cluster (BGC).^3^ There are many tools for identifying BGCs from genomes, including antiSMASH,^4^ PRISM,^5^ DeepBGC,^6^ SMURF,^7^ GECCO,^8^ RODEO,^9^ SanntiS,^10^ BAGEL,^11^ DeepRiPP,^12^ NeurRiPP,^13^ decRiPPter.^14^ Once a BGC has been identified, it can be compared to a set of known BGCs to determine how likely that BGC is to produce a novel compound. There are many methods for comparing BGCs including knownclusterblast,^4^ MulitGeneBLAST,^15^ CAGECAT,^16^ BiG-SCAPE,^17^ BiG-SLiCE,^18^ clust-o-matic,^19^ and lsaBGC.^20^ However, structural novelty alone is not enough to lead to the discovery of a new lead compound – the natural product must also have bioactivity that is relevant to a human disease. Recently, several tools have been reported to address this issue by predicting bioactivity from information contained in the BGC including DeepBGC,^6^ PRISM 4,^5^ and a method we developed for predicting natural product bioactivity.^21^ These methods are severely limited by a lack of training data. For example, our method relies on a training set that was built from the MIBiG database, the gold standard database in the field for linking BGCs to natural products, which is built from extensive collaborative community effort.^22^ At the time of the dataset creation, MIBiG lacked any bioactivity data, so bioactivity classifications had to be manually extracted from the literature, which was extremely time consuming and led to a dataset of 1003 BGC-activity pairs.^21^ There are likely many natural products with known BGCs and bioactivity that are not included in MIBiG and therefore are not included in our dataset. In addition, new natural products are regularly reported, and if the authors of these reports do not deposit their data to MIBiG, they instead need to be captured by the large community efforts that are undertaken to update MIBiG. If the process of monitoring the literature for new reports of natural products and their BGCs or bioactivity could even be partially automated, it would save significant time and enable more accurate machine learning models for a variety of applications in natural product research.

There has long been an interest in automated extraction of data from publications and other forms of text, but the utility of these approaches was limited by low accuracy and lack of ability to deal with data in varying formats.^23^ Recently there has been a rapid advancement in large language models (LLMs) ability to interpret and generate text^24-26^ as well as multimodal deep neural networks that can work with multiple media forms such as text and images.^27,28^ One such model is ChatGPT-4o, offered as a chatbot interface by Open AI.^29^ There are several examples of ChatGPT and other language models being used to extract data from the literature across multiple fields of science including materials science,^30-32^ biomedical research fields,^33-35^ and general science applications.^36-38^ ChatGPT has also been investigated as a tool to extract data from medical notes.^39^ Beyond applications in automated data extraction, ChatGPT and other LLMs have also found use in various chemistry applications including answering questions about molecular properties or reaction yields,^40^ and even planning and performing experiments.^41,42^ LLMs have the potential to speed up the expansion of training datasets for natural product bioactivity prediction and other tasks in natural products research.

To test the ability of ChatGPT to extract data for natural product bioactivity prediction, we tested ChatGPT’s ability to extract the data needed to expand our training data set from the PDF file version of publications. This information includes noting whether the paper describes the discovery of a natural product, what the name of the natural product discovered is, if the paper reports bioactivity and if so, what was discovered about the bioactivity, if the paper describes the BGC and if so if it reports an accession number for the BGC or for the genome. We then compared these results to manually curated annotations to determine how accurate ChatGPT is at extracting this information and where common failures occur.

## Results and Discussion

### Curation of benchmark data

We initially set out to assemble a dataset of papers to serve as a benchmark to assess ChatGPT-4o’s ability to accurately extract data from the PDF version of a publication. To this end, we manually analyzed each paper for the following information – whether the paper reported the discovery of a natural product, what natural products were described in the paper, if the activity was reported and if so what was reported about the product’s activity, if the BGC was described and if so if there was a accession number for either the BGC or the genome (Supplementary Data Table 1). When assembling this dataset, we focused mostly on papers related to natural product discovery or those that contain descriptions of their biosynthesis or bioactivities. However, we made sure to select some papers more distantly related to natural products, such as papers that described the total synthesis or derivatization of natural products to determine whether ChatGPT-4o frequently hallucinates information about natural products that is not reported in these types of papers such as bioactivity or BGC.

### ChatGPT can successfully extract data on molecular names, bioactivity, and biosynthetic gene clusters from publications on natural products

We downloaded a PDF for each paper in our benchmark set. We then uploaded that paper to ChatGPT-4o and asked it a series of questions, following a workflow illustrated in Figure 1. We designed our series of prompts in order to avoid hallucinations, or when a model outputs incorrect or even nonsensical information, which is a common problem for LLMs.^43^ Instead of directly asking ChatGPT for information contained in the paper, we first asked a yes or no question to determine if the information was present in the paper at all. If ChatGPT responded “yes,” we then proceeded to ask for the information. We then scored each question we asked ChatGPT based on if the output matched our manually assigned description of the paper (Figure 2). Summaries of ChatGPT answers to questions and our scoring of these answers is available in Supplementary data table 2-11. The exact text used in the prompts is provided in the methods section.

**Figure 1.**
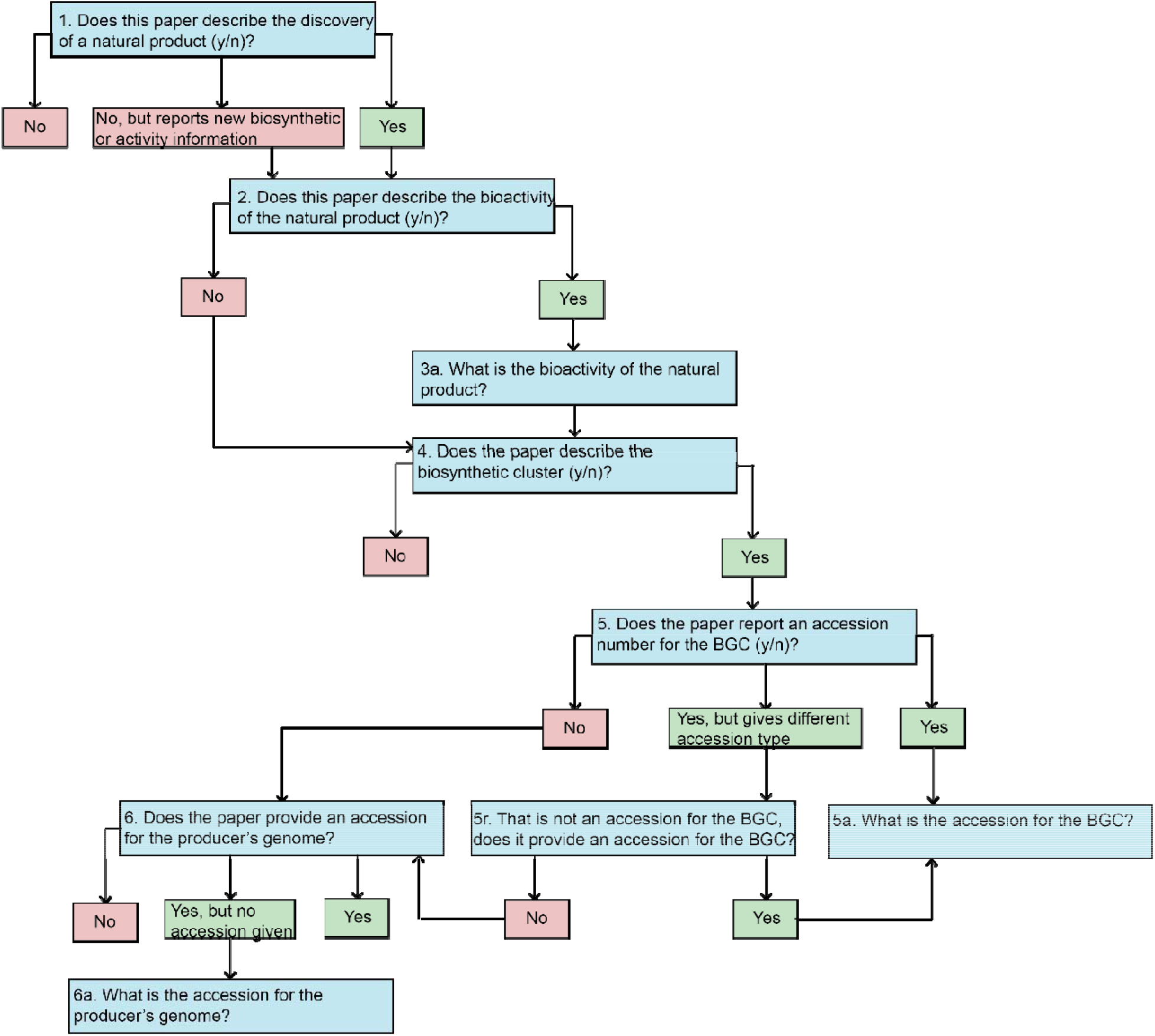
Prompting Workflow. This flow chart illustrates the prompting workflow we used to extract data with ChatGPT-4o. Questions asked by the prompter are shown in blue and ChatGPT responses we made prompting decisions from are shown in red or green.

**Figure 2.**
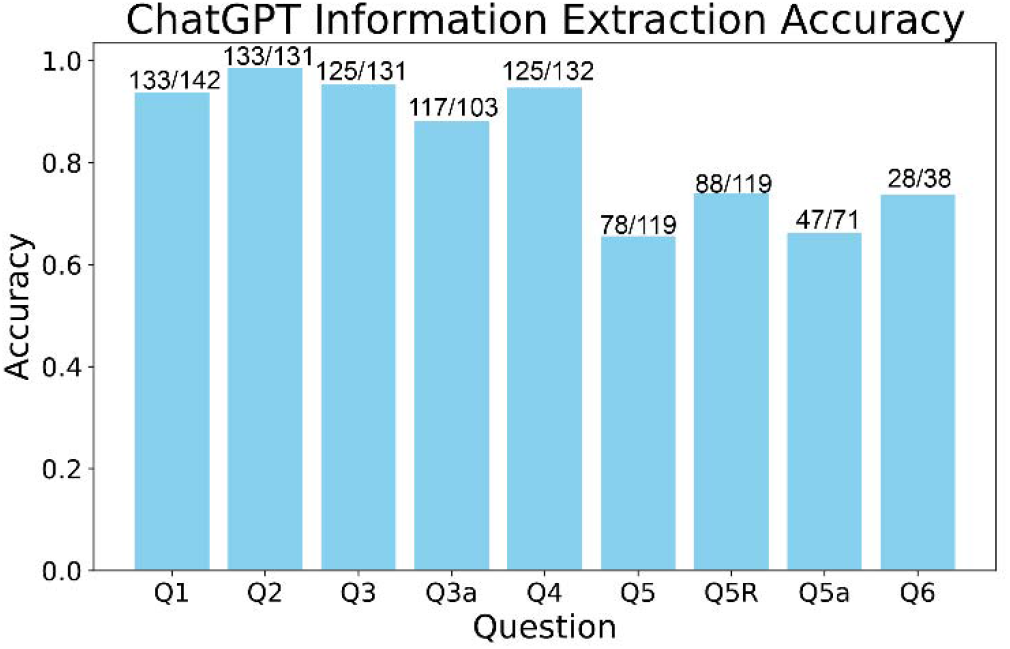
ChatGPT Accuracy. This bar graph shows the fraction of questions answered accuracy for each question asked. Numbers above the bars indicate the number of correct answers over the total number of questions. Questions 5-6 related to accession numbers show lower accuracy than questions 2-4 related to the natural product name and activity.

We first asked ChatGPT if the paper described the discovery of a new natural product. ChatGPT performed well on this question, answering 133 out of 142 questions correctly (94%). The most frequent mistake (eight examples) occurred when ChatGPT described a study that followed up on a previous report of a natural product to further study its structure, biosynthesis, activity, or that rediscovered a known compound from a previously unknown producer as a novel discovery. But since the goal of our work is to extract BGC-natural product-activity relationships, this type of error does not significantly affect our downstream work. In one case, ChatGPT described a study that used chemoenzymatic synthesis to generate non-natural fluorinated analogs of a natural product^44^ as a paper that had discovered a new natural product. ChatGPT did however note that these compounds were “new-to-nature”, indicating that the derivatives are not in fact naturally occurring compounds. ChatGPT did well at recognizing papers that described the total synthesis or synthetic derivation of natural products and correctly characterized all seven of these papers as not describing a novel natural product.

For the next question, we asked ChatGPT to name the natural products discovered in the paper. ChatGPT also performed well at this task, listing the correct novel natural products without for 131 out of 133 questions (98%) correctly. In one case, ChatGPT hallucinated the names of natural products reported in the paper. In that instance, the paper reported the discovery of microviridin N9, but ChatGPT instead answered that the paper reported nostocyclopeptide M1 and M2 and nostopeptolide. Nostopepolide was mentioned in the paper, although it was not the main focus. Nostocyclopeptide M1 and M2 were only mentioned in a title in one of the references.^45^ In another case, ChatGPT identified a natural product discussed in the paper, but did not list the novel product.

We next asked ChatGPT a yes or no question about if the paper reported bioactivity for the natural products described in the paper. ChatGPT performed well on this task as well, answering 125 out of 131 (95%) correctly. In one case, ChatGPT answered that the paper described the bioactivity of the natural product when the paper did not; but within the response, ChatGPT also specified that it was referring to a different set of natural products, rhamnolipids, that was found to be an elicitor of the natural products, the albubactins, discovered in the paper. ChatGPT’s response to the question, “Yes, the paper describes the bioactivity of the albubactins. The study demonstrates that rhamnolipids are effective at significantly increasing the secondary metabolite production of the new lipopeptides albubactins A−H and albubactin acids A−C,” showed this clear contradiction as ChatGPT is clearly referring to the activity of the rhamnolipids not the albubactins. The paper does report that rhamnolipids induce production of the albubactins,^46^ so this information is not hallucinated but rather misinterpreted by ChatGPT. In two cases, ChatGPT answered that the paper reported activity of a natural product but then went on to describe activity of one of the enzymes involved in the production of that natural product or the product’s biosynthesis. ChatGPT also sometimes does not correctly process brief references to the bioactivity present in the introduction of papers. For example, one paper states, “Because SapB lacks apparent antibiotic activity, the term lantipeptide has been coined to describe it”^47^ but ChatGPT states “No, the paper does not describe the bioactivity of the compound SapB,” even though the paper clearly states that SapB is known to not have antibacterial activity. In one case, ChatGPT initially answered that the paper did describe the bioactivity, but in the next follow up question admitted that the paper did not actually contain that information.

If ChatGPT answered yes to question 3, we then asked a follow-up question (3a) where we prompted ChatGPT to describe the activity reported in the paper. ChatGPT did not perform as well on this question compared to the previous questions, answering 103 out of 117 (88%) correctly. In most of the cases where ChatGPT was incorrect (nine answers), ChatGPT identified some of the information reported in the paper correctly but missed other pieces of information. ChatGPT twice described the activity of an enzyme involved in biosynthesis rather than activity of the natural product, and in one other case, ChatGPT listed the classes of the product described rather than their activities. In one case, ChatGPT reported a technically correct but ambiguous answer, reporting that a compound described as having antibacterial activity in the original paper had antimicrobial activity. There was also a case where this question resulted in a correction to the answer reported in question 3. For one of the papers, ChatGPT responded to question 3 to state that the paper reported information about the bioactivity even though it did not.^48^ ChatGPT then gave the following answer to question 3a “The paper does not specifically detail the bioactivities of Tambjamine BE-18591 in the provided excerpts. However, it mentions that tambjamines, in general, display a wide array of bioactivities, including antibacterial, antimalarial, immunosuppressive, and antitumor activities. For precise details on the bioactivity of Tambjamine BE-18591, you may need to refer to the specific sections of the paper dedicated to its bioactivity.”

Regardless of the answer to question 3, we asked ChatGPT if the paper described the BGC of the natural product. ChatGPT performed well on this question, answering 125 out of 132 (95%) correctly. In the majority (five) of cases where ChatGPT answered this question incorrectly, the paper did describe at least one gene involved in biosynthesis and sometimes also described efforts to clone the BGC without giving specifics of the BGC itself. In one of the remaining incorrect examples, ChatGPT mistook discussion of a regulator of biosynthesis that was manipulated in the paper^49^ for a discussion of the BGC. This was immediately apparent from the text of the response “Yes, the paper describes the biosynthetic gene cluster that produces these compounds. It mentions the identification and expression of a previously uncharacterized SARP protein, Syo_1.56, in Streptomyces sp. RK18-A0406, which led to the discovery of the elasnin.” For the final incorrect answer, the paper did not discuss the biosynthesis of the compounds at all, and it is not clear why ChatGPT responded that the paper did discuss the BGC.

### ChatGPT has significantly lower accuracy extracting the correct accession numbers

Next, we investigated if ChatGPT could extract accession numbers for the BGCs for papers that ChatGPT reported as describing a BGC. To accomplish this, we first asked a yes or no question about if an NCBI accession for the biosynthetic cluster was reported. During our initial testing, we found that ChatGPT often answered with more detail than a yes/no response and that this detail often revealed that ChatGPT was responding with a different type of accession number - for example, the accession for the full genome sequence or an accession number for NMR data. Therefore, we decided to add an optional “rescue” prompt after the accession question that the user would ask if ChatGPT revealed that it was responding with the wrong type of accession. ChatGPT answered the first question correctly only 78 out of 119 times (66%). The rescue prompt was used for ten of these incorrect answers, and ChatGPT successfully recognized that it had provided the wrong type of accession and admitted that the paper did not have the accession for the BGC in all ten cases, bringing the overall correct response rate up to 74% after rescue. For every incorrect answer, ChatGPT answered “yes” when the correct answer was “no”. In some cases, ChatGPT would suggest that the user check the supporting information for the accession, despite the paper not mentioning that accessions were provided in the supporting information. This suggests that ChatGPT could be hallucinating this information due to a prevalence of suggestions to check the supporting information for additional information in its training set.

If ChatGPT responded that the BGC accession was in the paper, we then asked it to provide the accession number. ChatGPT also struggled with this question, answering 47 out of 71 (66%) of questions correctly. In three cases, ChatGPT corrected its previously incorrect answer, admitting that there was no BGC accession reported in the paper. The most common error, which occurred 19 times, was that ChatGPT reported a correct accession for either individual genes or the producer’s whole genome rather than an accession for the BGC. In four of these cases, the text response noted that the accession was for something other than the BGC. In the remaining cases, ChatGPT reported the accession and stated that the accession was for the BGC. This implies that ChatGPT has difficulty determining types of accessions, even when the text of the paper clearly reports what the accession number is for. ChatGPT also sometimes hallucinated a response. This occurred three times, with ChatGPT reporting a real accession number that was not present in the paper and was not at all related to the contents of the paper. For example, in one case ChatGPT reported the accession number for a bronchitis virus strain despite bronchitis not being mentioned in the paper. This suggests that ChatGPT memorized these accessions during its training and sometimes reports them despite their lack of relation to the paper. However, despite the general inaccuracies, ChatGPT did report the correct accession whenever that accession was described in the paper in all but one case. In that case, the paper was an older PDF that did not have modern text encoding and ChatGPT mistook a “Z” for a “2” in the accession number.

If ChatGPT responded that the BGC accession was not in the paper, we then asked if the paper contained the accession for the producer’s genome. In response to the yes or no question, ChatGPT answered correctly 28 out of 38 times (74%). Generally, ChatGPT provided the accession along with the yes or no answer, but if it did not, we further queried ChatGPT to ask what the accession was. In two cases, this resulted in ChatGPT correcting itself to admit that there was no accession reported in the paper. As with question 5, most inaccuracies were due to ChatGPT reporting the incorrect type of accession such as the accession for the organism’s 16S sequence, accession numbers for individual genes, or a link to a database where the paper obtained information from but without an accession. ChatGPT also hallucinated on this question and twice reported accession numbers for genomes of bacteria unrelated to the producer. Interestingly, in two cases ChatGPT reported the accession for the same species described in the paper, even though the paper did not mention the accession number, which suggests that ChatGPT learned the association between the accession and the species from its training set. As with the previous question, when a genome accession was available in the paper, ChatGPT always reported the correct accession and it was only incorrect in cases where no accession was available.

### An altered prompting strategy can improve accuracy

We wondered if an altered prompting strategy could improve the accuracy of some of the less accurate questions. Crafting of prompting strategies, also known as “prompt engineering” has been shown to improve accuracy.^50^ We tried replacing question 5 with the question “Does the paper report an accession number for the biosynthetic cluster that produces the compounds(s), yes or no? Be sure to only consider accessions for biosynthetic clusters and not other types of accession, including the accession for genome sequences.” We hypothesized that this question would prevent some of the cases where ChatGPT reported a genome or other accession instead of a BGC. We reran the new prompting workflow on 31 of the papers for which ChatGPT reported a non-BGC accession as a BGC accession. With the altered prompting strategy, ChatGPT now gave the correct response for 23 of those 31 papers. This result demonstrates that changes made in prompting strategies designed to avoid common errors can greatly improve accuracy of responses.

## Conclusion

Assembly of robust datasets related to natural product bioactivity and biosynthesis is essential for development of machine learning and artificial intelligence models related to natural products. Historically, the field has relied on extensive community efforts to mine the literature, but there is likely still a lot of data in the literature that is not included in existing databases. In this work, we investigated the ability of ChatGPT to extract data from papers about natural products. We found that ChatGPT performed well at determining if a paper described a novel natural product, reporting the name of the product, describing the bioactivity of the product, and identifying if the paper described the BGC with accuracies ranging from 94 to 98%. ChatGPT was less accurate at extracting accession numbers for BGCs or producers’ genomes, with accuracies ranging from 65 to 74% on questions related to that topic. In particular, ChatGPT reported the wrong type of accession or an accession that was not present in the paper. We found that altering the prompting strategy to instruct ChatGPT to not report the accession other than the one of interest could improve accuracy. With further improvements to the technology underlying ChatGPT and prompting strategies, ChatGPT will likely soon reach the required accuracy to fully automate the mining of the literature for data on natural products. This will lead to significant improvements in emerging machine learning and artificial intelligence models used in natural product discovery.

## Methods

### Assembly of benchmark data

The papers used to create the benchmark dataset in this study were originally assembled by various lab members in the course of their respective research. Papers were identified both from existing databases such as MIBiG as well as simple web searches. All papers come from peer reviewed journals or preprint servers and were manually annotated by the authors.

### Prompting strategy

1. (Upload pdf) Does the attached paper describe the discovery of a novel natural product, yes or no?
2. What are the names of the natural products discovered in this paper?
3. Does the paper describe the bioactivity of these compound(s), yes or no?
  a. (if yes to 3) What is reported about the bioactivities of the compounds described in the paper?
4. Does the paper describe the biosynthetic gene cluster that produces these compounds(s), yes or no?
5. Does the paper report an accession number for the biosynthetic cluster that produces the compounds(s), yes or no?
  a. What is the NCBI accession for the BGC?
  r. (if it gives NMR or other accession to 6, use this with <> replaced with the type of accession) That is an accession for the <NMR, MassIVE, GNPS data>, not the biosynthetic cluster sequence. Does it report an accession for the cluster?
6. (if yes to BGC and no to BGC accession and it hasn’t already provided the genome accession) Does the paper provide an accession for the producer’s genome?
  a. (if it hasn’t already provided the number in 6) What is the accession for the producers genome?

### Testing of prompt strategy

Questions 1, 3, 4, 5, 6 were yes or no questions. We scored ChatGPT as correct only if it gave the correct yes or no response, regardless of what other information was given in the answer that may have contradicted the yes or no.

For question 2, we scored the model as correct if it replied with the correct name of the all the novel products described in the paper. If a model reported additional compounds that were also discussed in the paper, we did not count that as incorrect as long as the model reported the main novel compounds as well. However, if the model reported a product that was not discussed at all in the paper, we scored it as incorrect.

For question 3a, we scored ChatGPT as correct if it identified the same types of activity or inactivities as were described in the paper. If ChatGPT missed one of the major activities described in the paper, described an activity that was not described in the paper, or described activity unrelated to the bioactivity of the natural product described in the paper, we scored it as incorrect. If ChatGPT gave a response that admitted that its previous response to question 3 was incorrect and the paper did not actually describe bioactivity, we scored it as correct. If ChatGPT gave a technically correct answer that was too ambiguous for our downstream applications, we also scored it as incorrect. For example, if ChatGPT reported that a compound had antimicrobial activity, but the paper reported a more specific label such as antibacterial or antifungal, we marked that as incorrect. If there were multiple compounds described in the paper, we only scored ChatGPT based on its response for the compounds that were the main focus of the paper.

For questions 5a and 6a, matches were determined based on direct comparison with the benchmark set. If ChatGPT offered other accessions, even for individual genes within the same genome, they were scored incorrectly. If for question 5 or 6 ChatGPT erroneously reported an accession existed when there was none and then admitted that in 5a or 6a, question 5a or 6a was scored as correct.

### Altered prompting strategy

We replaced question 5 with the following prompt:

Does the paper report an accession number for the biosynthetic cluster that produces the compounds(s), yes or no? Be sure to only consider accessions for biosynthetic clusters and not other types of accession, including the accession for genome sequences.

And reran the entire prompting strategy up to question 5 for 31 of the papers ChatGPT gave an incorrect response to question 5 and that did not provide enough information in that response for the prompter to recognize that the accession was incorrect.

## Data and Software Availability

All manual annotations of publications are available in Supplementary Table 1. Original papers are available from their publishers, and we have provided the doi for each paper in Supplementary Table 1 so that they can easily be searched for. All summaries of ChatGPT output and our scoring is also available in the supplementary data tables. No original software or code was created for this publication. ChatGPT-4o is available on a limited basis for a free account or purchase through a subscription from OpenAI: https://openai.com/chatgpt/

## Acknowledgements

This work was supported by the National Institutes of Health, grant number R35GM146987. The content is solely the responsibility of the authors and does not necessarily represent the official views of the National Institutes of Health.

## Table of Contents Figure

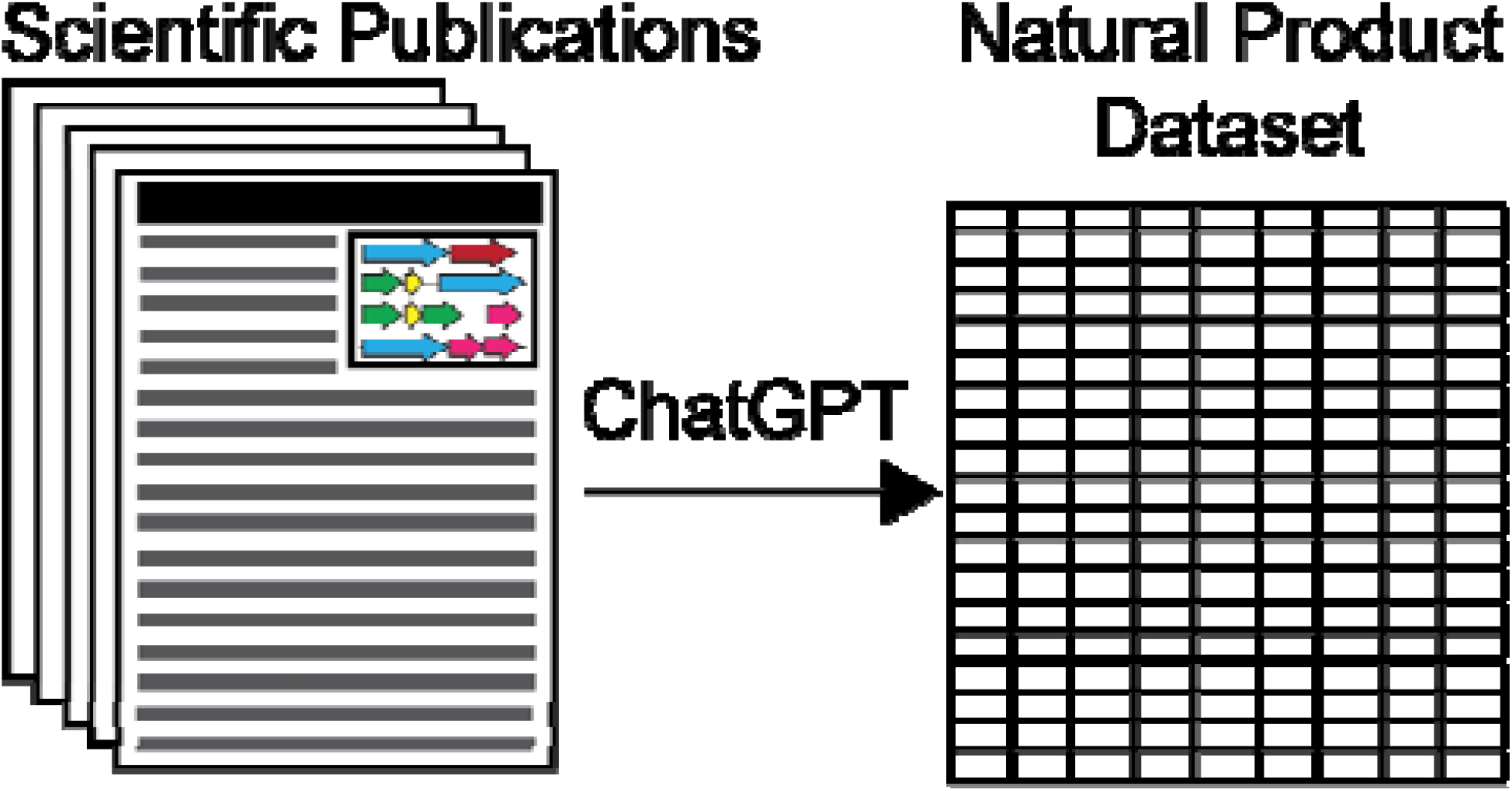

